# Deconstructing natural products to develop synthetic small molecule attachment inhibitors with broad spectrum antiviral activity

**DOI:** 10.1101/2025.05.20.655152

**Authors:** Consuelo C Correa-Sierra, Jody L Cameron, Seyedeh Nargess Hosseini, Devon Schatz, James Donnelly, Chloe Murrell, Furkat Mukhtarov, Che C Colpitts, Frederick West, Luis M Schang

## Abstract

The continuous emergence of new viruses and the number of viruses that are each highly consequential for few people raise a need for broad spectrum antivirals. Most human and emerging viruses first attach to cellular glycosaminoglycans (GAG) or sialylated glycans (SG), a potential target for broad spectrum antivirals. Attachment to the former is through polar interactions between the negatively charged glycans and positively charged domains in virion proteins and typically inhibited by negatively charged polymers. Attachment to the latter is through specific interactions at binding pockets and inhibited by molecules that bind to these pockets. Surprisingly, EGCG inhibits viruses that attach to GAG and SG. However, it does so with widely differing potencies, is not a pharmacologically desirable molecule, and is limited by solubility. We tested whether it was possible to develop small synthetic molecules to inhibit viruses that attach to SG or GAG. We first identified the EGCG moieties responsible for the antiviral activity. The two polyhydroxylated phenyl groups were essential while the central benzopyran linker was not. We thus designed a series of gallate compounds to explore the minimal pharmacophore required for broad-spectrum antiviral activity. By exploring the linkers and number of galloyls, we identified small molecule inhibitors of herpes simplex virus 1, influenza A virus, and the coronaviruses hCoV OC43 and SARS-CoV-2. These compounds have low micromolar to submicromolar potency and no limiting cytotoxicity. These molecules are still not pharmacologically optimized and limited by solubility, but they define a minimal pharmacophore that confers broad spectrum antiviral activity.

**Importance:** The SARS-CoV-2 pandemic demonstrated the importance of antivirals in managing emerging viruses. Although vaccines were successfully developed in less than two years, there was resistance to vaccination while infected or sick people were far more willing to take antivirals. It is impossible to develop antivirals specific for unknown viruses, but broad spectrum antivirals could control viral spread until more specific and potent drugs are developed. Human pathogenic and emerging viruses commonly attach to glycans, providing a target for broad spectrum antivirals. However, inhibitors of attachment to glycosaminoglycans do not typically inhibit viruses attaching to sialylated glycans, and vice-versa. We had found that EGCG has the unique property of inhibiting viruses that attach both glycans, with quite different potencies. Here, deconstructed a natural compound, EGCG, to identify the moieties responsible for its antiviral activity to then produce broad spectrum antiviral compounds against established and emerging viruses that attach to either glycan.

## Introduction

The SARS-CoV-2 pandemic highlighted the continuous threat posed by emerging viruses and the challenges in managing them. Although vaccines were successfully developed in less than two years (1), healthy people were often unwilling to be vaccinated (2). In contrast, infected or sick patients were less resistant to therapeutic (3), or even allegedly therapeutic (4, 5), treatments. This experience has shown that beyond vaccines, antivirals are critical to aid in minimizing the impact of a pandemic on health, lives, the economy, and society.

The development of antivirals against emerging or re-emerging viruses presents a difficult challenge and requires significant investment of unknown returns (6). Several approaches have been proposed to overcome this bottleneck, including general drug repurposing (7-12), natural products (13, 14), and broad spectrum antivirals (15-17).

The many unsuccessful attempts to repurpose drugs against SARS-CoV-2 (18, 19) highlight the challenges to drug repurposing. Several natural products have been developed as clinical drugs, including antibiotics, antiparasitic, and antivirals, but natural products are often limited by pharmacological properties or their amenability for chemical optimization. Broad spectrum antivirals targeting conserved viral proteins, such as RNA-dependent RNA polymerases (RdRP) were successfully repurposed against SARS-CoV-2 (15-17). Other potential targets for broad spectrum antivirals have been explored, including envelope lipids (20-28) and cellular proteins required for the replication of many unrelated viruses (29-38).

Glycans also play important roles in the replication cycle of many viruses. All but one established or known emerging human viruses, including the herpesviruses and SARS-CoV-2, attach to glycosaminoglycans (GAG) such as heparan sulfate, chondroitin sulfate, or hyaluronic acid (39). This attachment occurs via polar interactions between the negatively charged linear GAG and positively charged, often non-structured, domains in the viral surface proteins (39-43). Other viruses, like IAV or hCoV OC43 bind to a sialyl residue attached to a galactose at the termini of branched sialylated glycans (SG) via specific interactions at well-defined binding pockets (44-50). hCoV OC43 also binds to GAG (51), SARS-CoV-2 may also bind to SG (13, 52, 53), and there is one report of high concentrations of heparin derivatives inhibiting entry of viruses pseudotyped with IAV HA (54).

Binding to GAG is inhibited by negatively charged polymers (55-62), including biologicals and a number of natural products (62-68); these polymers do not typically inhibit attachment to SG. Binding to SG is inhibited by molecules presenting several properly oriented sialyl-mimetic structures (44, 67, 69-80), including natural products (67, 68, 75); these molecules do not typically inhibit attachment to GAG.

Glycan attachment is essential for viruses that attach only to them but not for those with secondary proteinaceous receptors. Nonetheless, increasing the number of virion binding sites for cellular GAG increases infectivity (81), binding increases attachment and infection by >10-100-fold (82), and mutants defective in glycan attachment are often structurally unstable and replication defective in culture, and attenuated in vivo (82-84).

Viral attachment is a validated antiviral target. Maraviroc inhibits HIV attachment to CCR5 (85), and several monoclonal antibodies that inhibit binding of spike to ACE2 were used against SARS-CoV-2 until they selected for resistance (86). However, no inhibitor of virion attachment to glycans has been developed as an antiviral yet. The major challenges for such development are that glycan attachment is often low affinity and cooperative, and that it proceeds through two different modes.

We and others had found that the natural catechin epigallocatechin gallate (EGCG) inhibits the attachment of HCV virions to cells (66). Considering the molecular shape and polarity of EGCG, we proposed that it could act as a competitive inhibitor for virion attachment to GAG. To test this model, we evaluated its activity against several viruses that attach to GAG, poliovirus, which does not attach to any glycan, and a virus that attaches to SG, IAV. EGCG inhibited the infectivity of all viruses that attach to GAG, but not that of poliovirus, as expected (67). Much to our surprise, EGCG also inhibited the infectivity of IAV, albeit with ∼50-100-fold lower potency than against viruses that attach to GAG. Mechanistically, EGCG inhibits binding of the virion proteins to GAG or SG, as shown by inhibition of HSV-1 attachment to cells or heparin, and of IAV attachment to cells and hemagglutination, respectively (67, 87). EGCG thus has the unusual ability to inhibit attachment of virions to the two different types of glycans.

Besides the large differences in relative potency against viruses that attach to GAG and SG (67), EGCG is not a pharmacologically desirable molecule, with poor PK/PD properties including a short half life (88-90). EGCG is not easily amenable to medicinal chemistry optimization and has different moieties, which could each inhibit separately viruses that attach to GAG or SG. We thus evaluated whether we could identify the moieties of EGCG critical for the inhibition of viruses that attach to GAG or SG to then synthesize simpler molecules and systematically evaluate their potency.

Here we report the identification of synthetic di- and poly-gallates with broad spectrum antiviral activity in the low to sub-micromolar range. Although these molecules contain pharmacologically suboptimal ester links and polyphenols, which confer metabolic vulnerability and limited solubility (91-96), they uncover the potential of targeting the conserved glycan attachment and binding step in broad spectrum antiviral drug development and identify a simple pharmacophore conferring this activity.

## Results

### Two polyhydroxylated phenyl rings are required to inhibit viral infectivity

The different moieties of EGCG could confer different antiviral activities against viruses binding to each type of glycans. We started testing for this possibility by evaluating the activities of different catechins against the infectivity of the GAG-binding HSV-1 and the SG-binding IAV. To compare the ability to inhibit the infectivity of both types of viruses, we developed the “broad spectrum range” (BSR), as the ratio of the highest to the lowest EC_50_, and the “broad spectrum index” (BSI), as the multiplication of the BSR by the highest EC_50_. For both factors, lower is better (Table 1). We were interested in identifying molecules targeting virion proteins but cells have complex surface glycocalyces which could well be targeted by the test compounds. We thus pre-treated the virions for 10 min at 37°C before infecting the cells to bias the identification towards compounds that act on the virions. Catechins with a galloyl and a second polyhydroxylated phenyl ring inhibited the infectivity of HSV-1 and IAV (ECG, Table 1 and Fig. 1), although with about 35-45-fold different potencies and high BSI (>200). Compounds with only one polyhydroxylated phenyl ring did not have observable antiviral activity (EC, Table 1 and Fig. 1). The benzopyran linker between the hydroxylated phenyl rings is not sufficient to confer any antiviral activity whereas the two polyhydroxylated phenyl rings are required to inhibit IAV and HSV-1, though the number of hydroxyl groups is not critical (EGCG vs ECG, Table 1 and Fig. 1). Neither of these catechins had obvious cytotoxic effects up to the highest concentrations tested, failing to affect cell doubling during 48 or 72 h of exposure to 100 µM (Table 2). The relative antiviral activities of the different catechins agree with those described by Gescher *et al.,* (87).

**Fig. 1.**
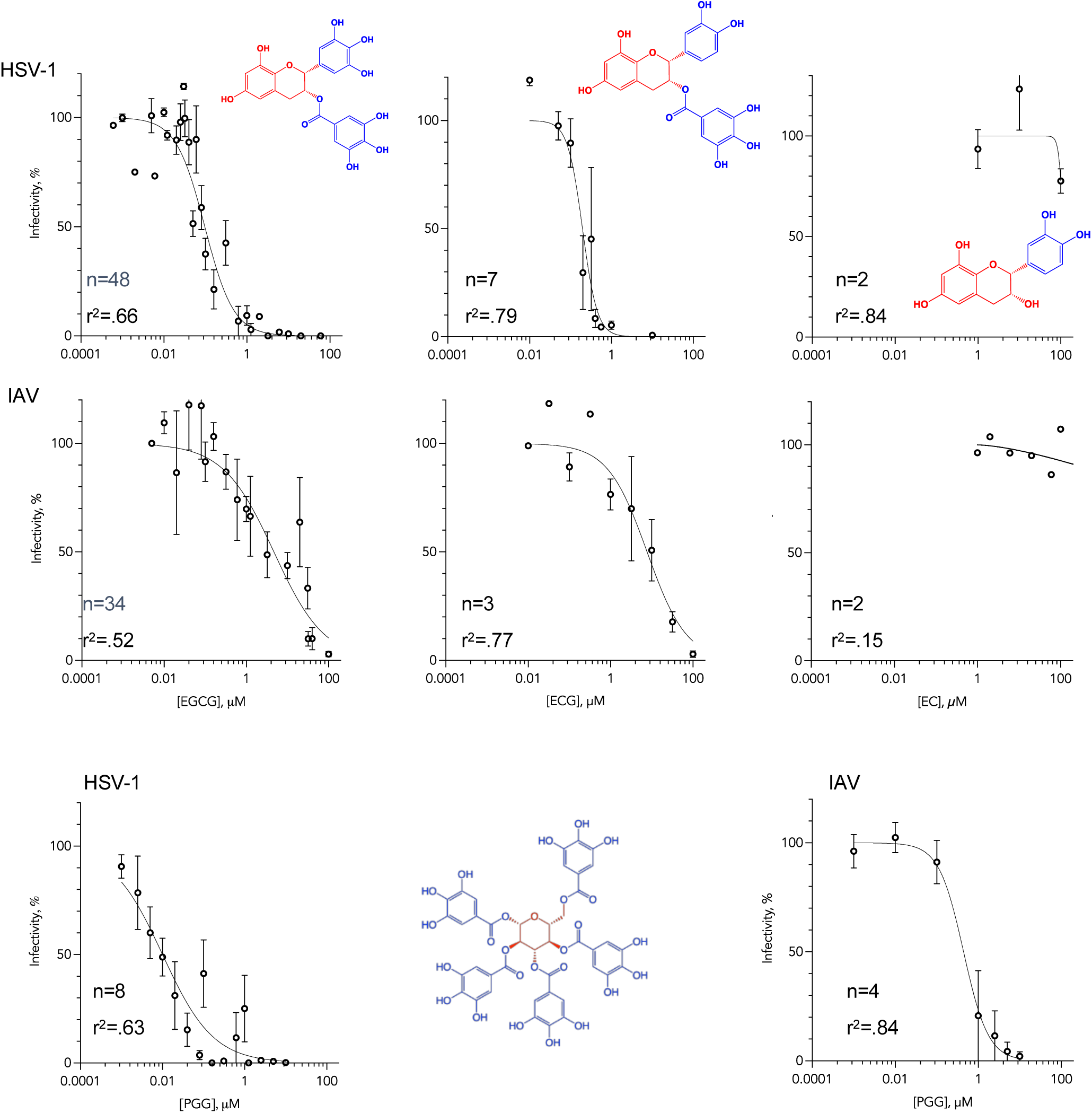
Inhibition of the infectivity of HSV-1 or IAV virions by different catechins and pentagalloyl glucose. Dose-response line graphs showing the inhibition of infectivity of HSV-1 or IAV by catechins or PGG. HSV-1 or IAV virions were mixed with the different catechins or PGG and incubated at 37°C for 10 minutes before infecting Vero (HSV-1) or MDCK (IAV) cells with 200 PFU. Average ± SEM, n= 23 to 48 biological independent experiments.

**Table 1.**
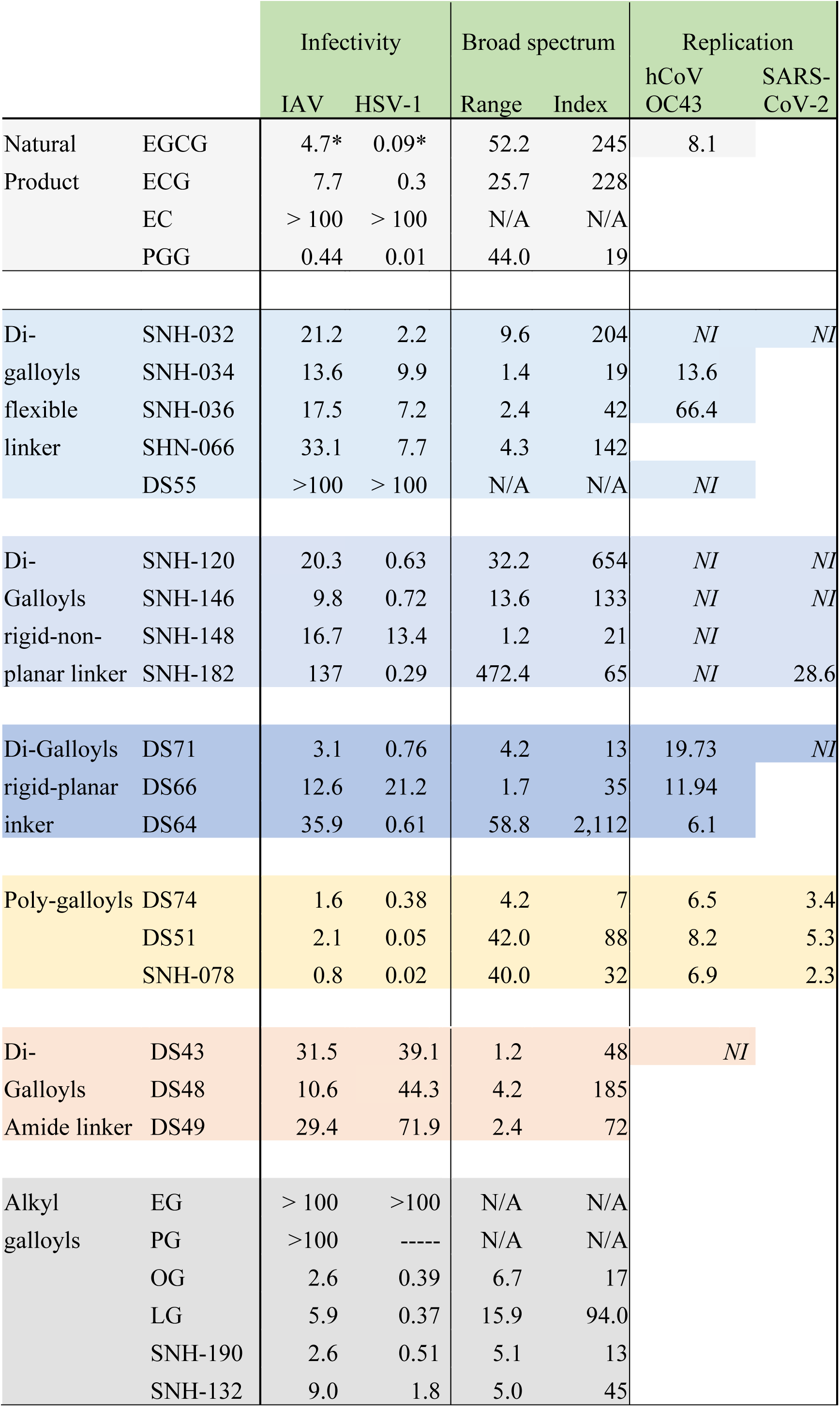
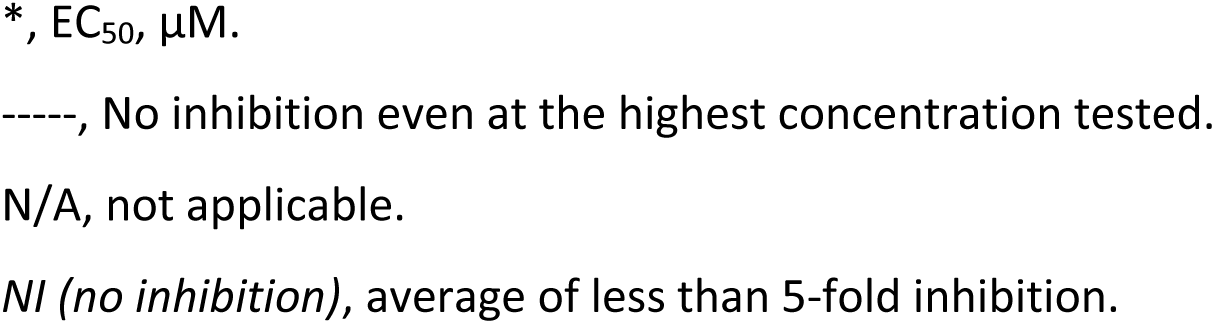
Antiviral potencies of the different galloys against HSV-1, IAV, OC43 and SARS-CoV-2.

|  |  | Infectivity |  | Broad spectrum |  | Replication |  |
| --- | --- | --- | --- | --- | --- | --- | --- |
|  |  | IAV | HSV-1 | Range | Index | hCoV-OC43 | SARS-CoV-2 |
| Natural | EGCG | 4.7* | 0.09* | 52.2 | 245 | 8.1 |  |
| Product | ECG | 7.7 | 0.3 | 25.7 | 228 |  |  |
|  | EC | > 100 | > 100 | N/A | N/A |  |  |
|  | PGG | 0.44 | 0.01 | 44.0 | 19 |  |  |
| Di- | SNH-032 | 21.2 | 2.2 | 9.6 | 204 | *NI* | *NI* |
| galloyls | SNH-034 | 13.6 | 9.9 | 1.4 | 19 | 13.6 |  |
| flexible | SNH-036 | 17.5 | 7.2 | 2.4 | 42 | 66.4 |  |
| linker | SHN-066 | 33.1 | 7.7 | 4.3 | 142 |  |  |
|  | DS55 | >100 | > 100 | N/A | N/A | *NI* |  |
| Di- | SNH-120 | 20.3 | 0.63 | 32.2 | 654 | *NI* | *NI* |
| Galloyls | SNH-146 | 9.8 | 0.72 | 13.6 | 133 | *NI* | *NI* |
| rigid-non- | SNH-148 | 16.7 | 13.4 | 1.2 | 21 | *NI* |  |
| planar linker | SNH-182 | 137 | 0.29 | 472.4 | 65 | *NI* | 28.6 |
| Di-Galloyls | DS71 | 3.1 | 0.76 | 4.2 | 13 | 19.73 | *NI* |
| rigid-planar | DS66 | 12.6 | 21.2 | 1.7 | 35 | 11.94 |  |
| inker | DS64 | 35.9 | 0.61 | 58.8 | 2,112 | 6.1 |  |
| Poly-galloyls | DS74 | 1.6 | 0.38 | 4.2 | 7 | 6.5 | 3.4 |
|  | DS51 | 2.1 | 0.05 | 42.0 | 88 | 8.2 | 5.3 |
|  | SNH-078 | 0.8 | 0.02 | 40.0 | 32 | 6.9 | 2.3 |
| Di- | DS43 | 31.5 | 39.1 | 1.2 | 48 | *NI* |  |
| Galloyls | DS48 | 10.6 | 44.3 | 4.2 | 185 |  |  |
| Amide linker | DS49 | 29.4 | 71.9 | 2.4 | 72 |  |  |
| Alkyl | EG | > 100 | >100 | N/A | N/A |  |  |
| galloyls | PG | >100 | ----- | N/A | N/A |  |  |
|  | OG | 2.6 | 0.39 | 6.7 | 17 |  |  |
|  | LG | 5.9 | 0.37 | 15.9 | 94.0 |  |  |
|  | SNH-190 | 2.6 | 0.51 | 5.1 | 13 |  |  |
|  | SNH-132 | 9.0 | 1.8 | 5.0 | 45 |  |  |

\*, EC<sub>50</sub>, µM.
-----, No inhibition even at the highest concentration tested.
N/A, not applicable.
*NI (no inhibition)*, average of less than 5-fold inhibition.

**Table 2.**
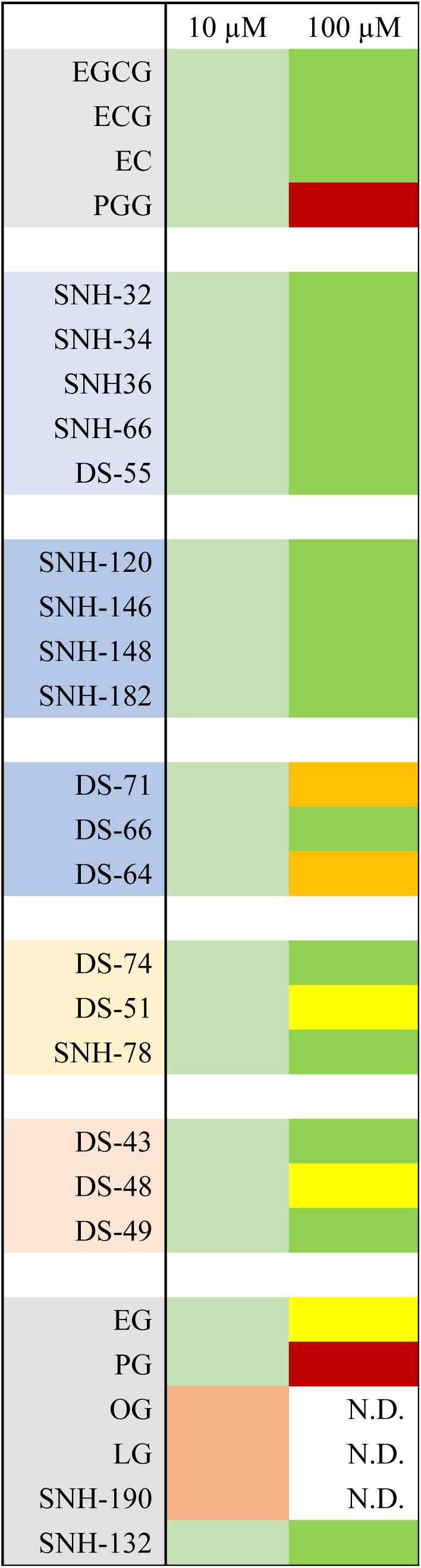
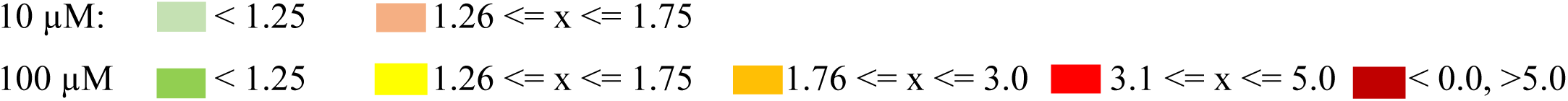
Analyses of cytotoxic and cytostatic effects. Subconfluent cells were exposed to 10 or 100 µM of the indicated compounds for 48 or 72 h and cell viability and doubling was evaluated by CellTiterGlo. No compound was cytotoxic (i.e. the number of viable cells continued to increase in the presence of all test compounds during the incubation) at 10 µM, but OG, LG and SNH190 increased cell doubling times. Only PGG and PG were cytotoxic (i.e., number of viable cells decreased in the presence of the compound) at 100 µM after 48 or 72 h, whereas some other compounds were modestly (i.e., doubling time increased by about 50 %, DS51 and DS48) or mildly (i.e., doubling times increased by about 2-fold, DS71, DS64) cytostatic. Orange, compounds that increased doubling time by ≥1.76-fold; dark red cytotoxic compounds. Compounds that were partially cytostatic at 10µM were not tested at 100µM (N.D.)

To further test the potential requirement for the benzopyran moiety, we tested another natural poly-galloyl, pentagalloyl glucose (PGG), which we and others have shown inhibits HCV (65). PGG has glucose instead of benzopyran as the linker between the galloyls. PGG inhibited the infectivity of HSV-1 and IAV with about 10-fold higher potency than EGCG or ECG, with a similar BSR of about 40, but a much lower BSI of almost 19, resulting from its much higher potency. PGG was cytotoxic after 48 h exposure to 100 µM, although it was not cytotoxic or cytostatic after 72 h exposure to 10 µM (Table 2), concentration at which it inhibited the infectivity of both HSV-1 and IAV by more than 99% (Fig.1). Attempts to test higher concentrations of PGG were unsuccessful, because of limited solubility.

### Digallate compounds inhibit the infectivity of GAG-HSV-1 and SG-attaching IAV virions

Based on the previous results, we proposed that at least two polyhydroxylated phenyl groups were required to inhibit the infectivity of viruses that attach to GAG or SG, that additional polyhydroxylated phenyl groups may increase potency, and that the benzopyran moiety was not required. To further test this conclusion, we synthesized a series of digallates with flexible alkyl linkers of different lengths. The rationale was that although the flexibility in the relative position of the putatively active galloyls would compromise binding affinity, and thus potency, it would allow us to evaluate the compounds without having to make any assumptions about preferred conformations.

All alkyl linked digallates inhibited the infectivity of GAG and SG-binding viruses regardless of linker length, but with only low to mid micromolar potency (Table 1 and Fig. 2). Although the BSR and BSI were generally lower than for the catechins and PGG, this was mostly a result of a decrease in potency against HSV-1, not an increase in potency against IAV. As expected for flexible linkers, there were only less than 2.5 or 5-fold differences in potency between compounds with linkers of 2 to 6 carbons against IAV or HSV-1, respectively. None of these compounds had obvious cytotoxic or cytostatic effects after 48 or 72 h exposure to 100 µM (Table 2). Addition of a polar oxygen atom in the linker disrupted antiviral activity against both HSV-1 and IAV (DS55, Table 1 and Fig. 2). Like the non-substituted flexible linkers, DS55 was not cytotoxic or cytostatic after 72 h exposure to up to 100 µM (Table 2).

**Fig. 2.**
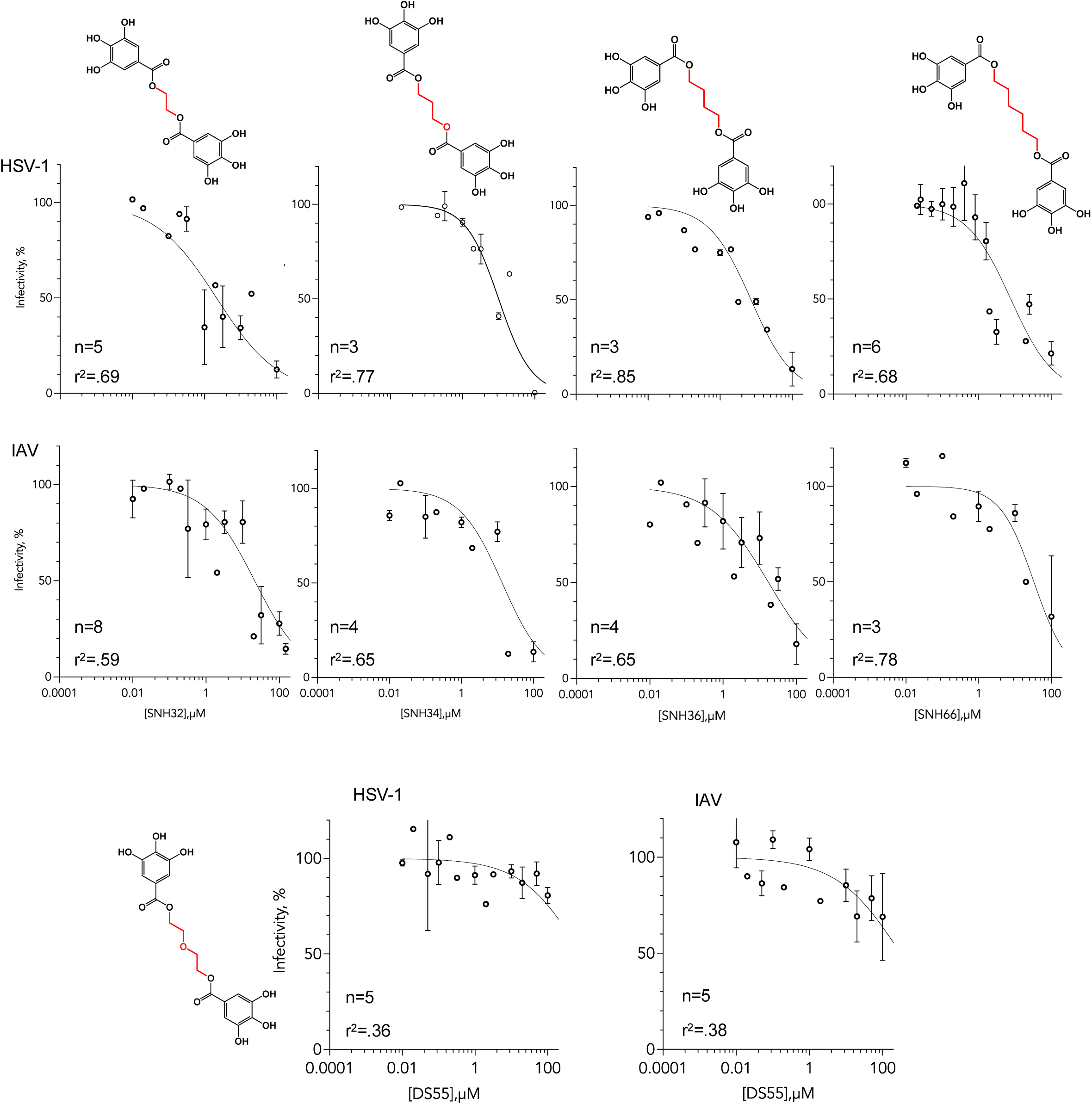
Inhibition of the infectivity of HSV-1 or IAV virions by digallates with flexible linkers. Dose-response line graphs showing the inhibition of infectivity of HSV-1 or IAV by the indicated digalates. HSV-1 or IAV virions were mixed with test compounds incubated at 37°C for 10 minutes before infecting Vero (HSV-1) or MDCK (IAV) cells with 200 PFU. Average ± SEM, n= 3 to 8 biological independent experiments.

### The conformation of the linker modulates antiviral potency

The linker in EGCG and PGG is rigid. We thus tested digallates with rigid linkers. Non-planar cyclohexane as a linker yielded compounds that inhibited the infectivity of the GAG binding HSV-1 with sub-micromolar potency and of the SG binding IAV at micromolar concentrations. For the cyclohexanes substituted at carbons 1 and 4, the *trans* isomer was the more active (Table 1 and Fig. 3). These compounds were consistently more potent against HSV-1 than IAV, but their BSR and BSI were higher than for the flexible linker compounds, except for SNH148 which had similarly low potency against both viruses (Table 1). Potency against HSV-1 was not influenced by the positions of the galloyls on the ring, with similar potency achieved for 1,4 or 1,2-disubstituted cyclohexanes (Table 1 and Fig. 3, SNH146 versus SNH182). However, the latter was mostly inactive against IAV, resulting in a BSI of more than 500,000. None of the compounds were cytotoxic or cytostatic after 72 h exposure to 100 µM (Table 2).

**Fig. 3.**
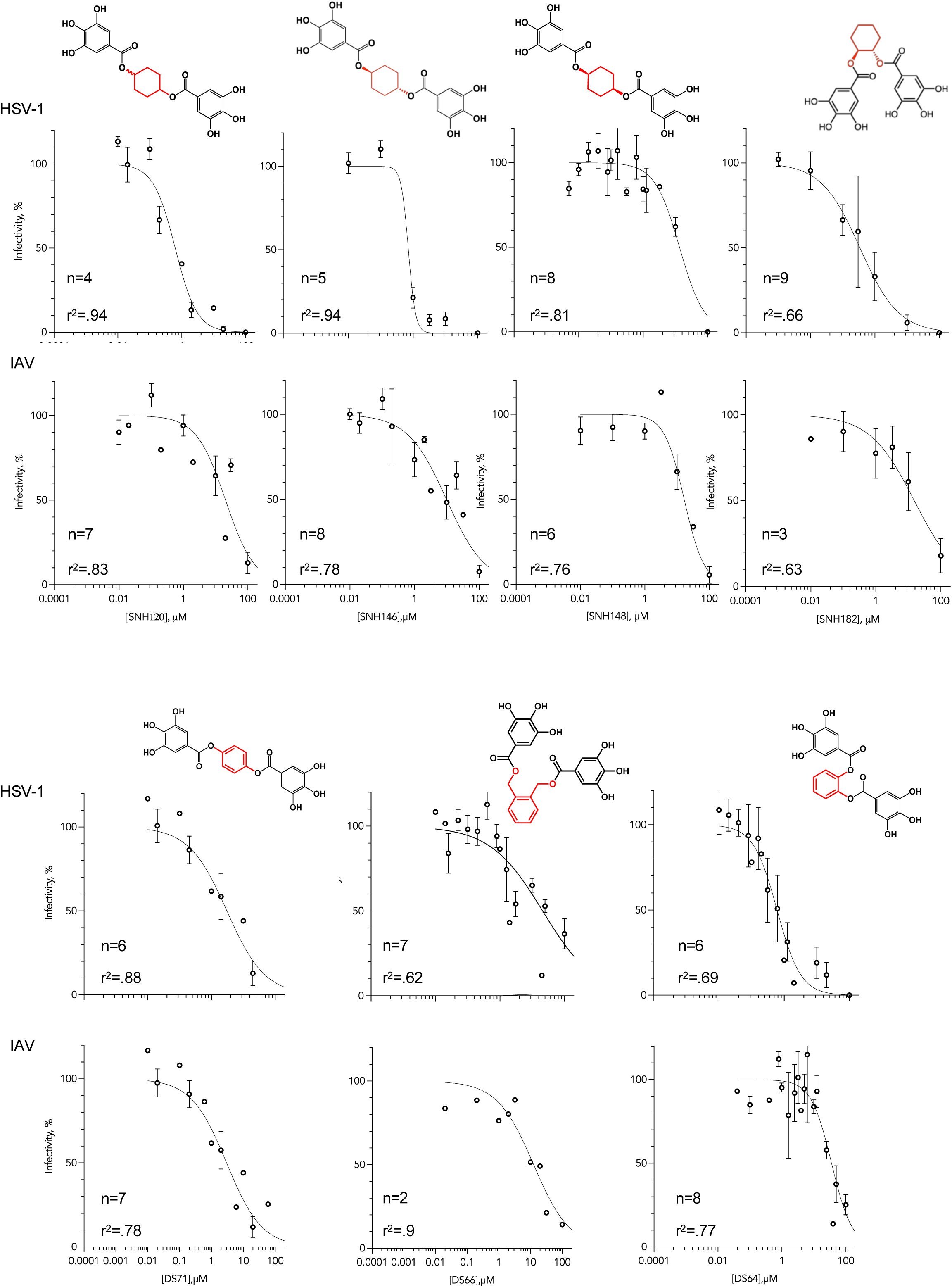
Inhibition of the infectivity of HSV-1 or IAV virions by digallates with rigid non-planar or planar linkers. Dose-response line graphs showing the inhibition of infectivity of HSV-1 or IAV by the indicated digallates. HSV-1 or IAV virions were mixed with test compounds incubated at 37°C for 10 minutes before infecting Vero (HSV-1) or MDCK (IAV) cells with 200 PFU. Average ± SEM, n= 2 to 9 biological independent experiments.

As the linker in EGCG is approximately planar and the relative position of the galloyls in PGG would be expected to be restricted by steric hindrance, we next tested a planar phenyl ring linker, which also yielded compounds with sub-micromolar to low micromolar potency. The substitution of the rigid non-planar by the rigid planar linker resulted in compounds maintaining the potency against HSV-1 while gaining potency against IAV (DS71 compared to SNH146 and SNH148 or DS64 compared to SNH182, Table 1 and Fig. 3). For compounds with polyphenols substituted para on the central benzene core (DS71 and DS66), the BSI was as low as 12.6, sigificantly lower than for the equivalent compound with a rigid non-planar linker (Table 1 and Fig. 3, DS71 compared to SNH146 and SNH148). The addition of flexible bonds connecting the polyphenyl groups to the rigid planar linkers disrupted antiviral activity against HSV-1, while slightly increasing the much weaker activity against IAV (DS64 versus DS66, Table 1 and Fig. 3). DS64 and DS71 were cytostatic, increasing cell doubling time by about 2.5-fold after 48 or 72 h exposure to 100 µM, but had no cytostatic or cytotoxic effects at 10 µM). DS66 had no obvious cytotoxic or cytostatic effects.

### Addition of a third or fourth galloyl yields more potent compounds

Considering that PGG was more potent than EGCG, and that addition of galloyl moieties commonly enhances biological activities of polyphenols (97), we next synthesized tri- and tetra-galloyl derivatives of flexible or rigid planar or non-planar linkers. This approach yielded potent compounds against both HSV-1 and IAV. The most potent, SNH78 (Table 1 and Fig. 4), inhibited the infectivity of GAG and SG-binding viruses with sub-micromolar potency, with a BSI of just 32. Although DS74 had the lowest BSI, at only 6.7, it was less potent against both viruses than SNH78. SNH78 and DS74 had no cytotoxic or cytostatic effects after 48 or 72 h exposure to 100 µM, whereas DS51 increased cell doubling time by about 1.5 fold. Attempts to test higher concentrations of the tetragalloyl SNH78 were unsuccesful, as were for the pentagalloyl PGG, because of limited solubility.

**Fig. 4.**
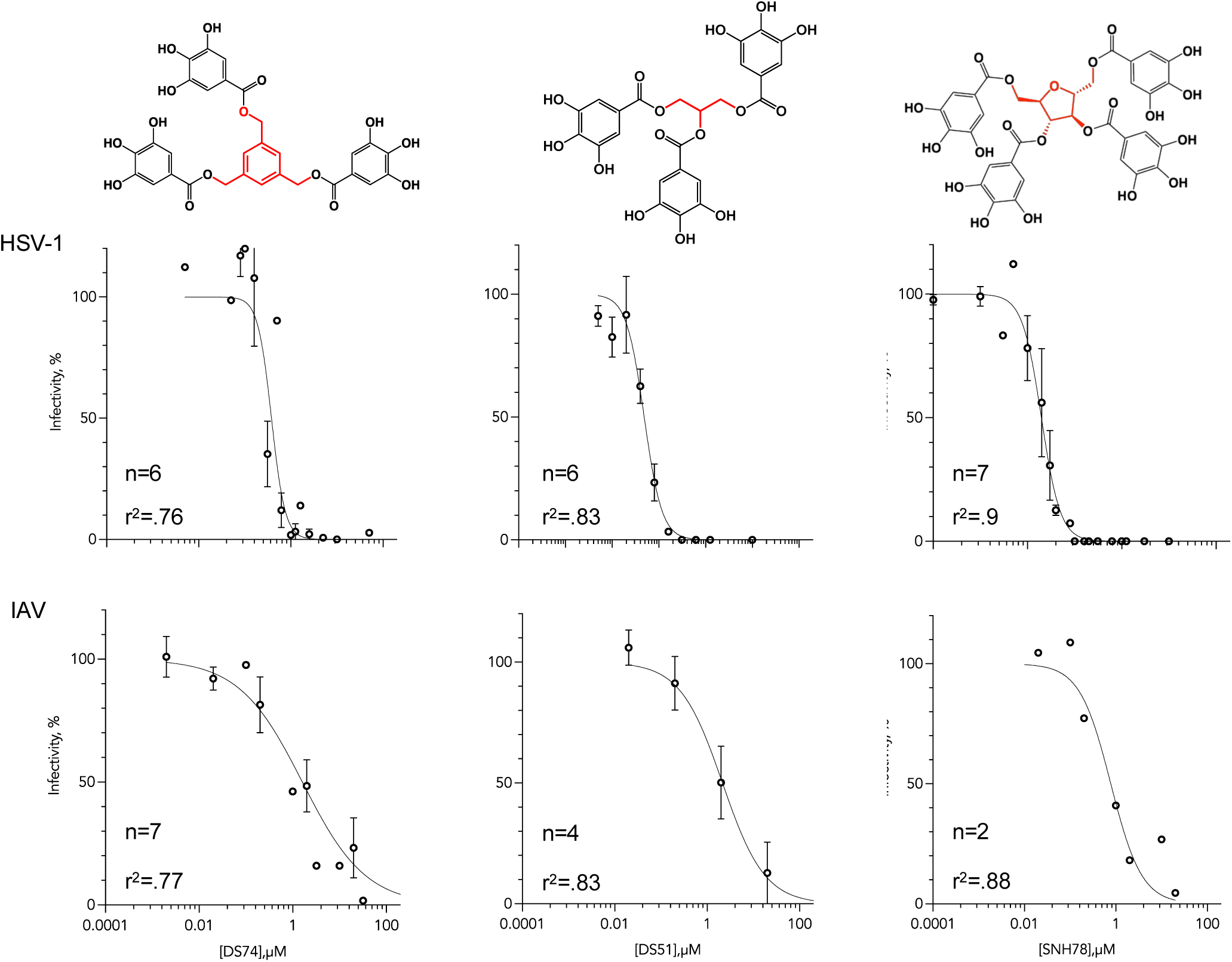
Inhibition of the infectivity of HSV-1 or IAV virions by synthetic polygalates. Dose-response line graphs showing the inhibition of infectivity of HSV-1 or IAV by the indicated polygalates. HSV-1 or IAV virions were mixed with test compounds incubated at 37°C for 10 minutes before infecting Vero (HSV-1) or MDCK (IAV) cells with 200 PFU. Average ± SEM, n= 2 to 7 biological independent experiments.

### Replacement of the esters by amide groups disrupts antiviral potency

Considering that ester links are not desirable in pharmacophores, due to their sensitivity to esterases (95, 96), we explored the replacement of the ester by amide groups. This replacement resulted in lower potency, particularly against HSV-1 (Table 1 and Fig. 5, DS43, 48, 49). DS43 or 49 did not have any cytotoxic or cytostatic effects after 48 or 72 h exposure to 100 µM, whereas DS48 was mildly cytostatic, increasing cell doubling times by 1.3 fold after 48 or 72 h exposure to 100 µM.

**Fig. 5.**
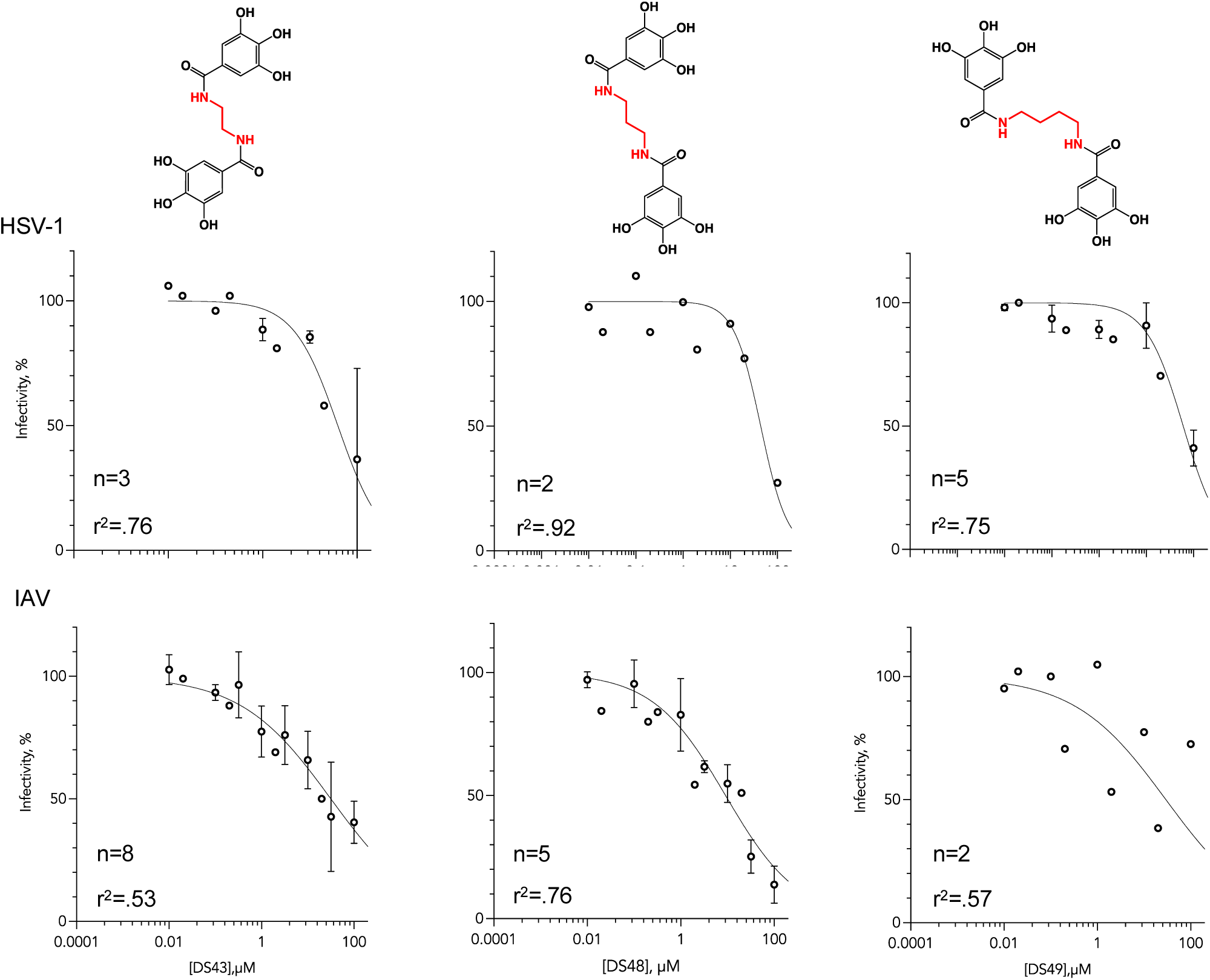
Effects of replacing the ester by amide linkers. Dose-response line graphs showing the inhibition of infectivity of HSV-1 or IAV by the indicated amide-linked polyphenols. HSV-1 or IAV virions were mixed with test compounds incubated at 37°C for 10 minutes before infecting Vero (HSV-1) or MDCK (IAV) cells with 200 PFU. Average ± SEM, n= 2 to 8 biological independent experiments.

### The ester link is not required for antiviral activity

We next tested the requirement for the ester group itself. To simplify the syntheses, we explored whether alkylgalloyls would inhibit the infectivity of IAV or HSV-1. Gallate esters of short-chain alcohols, ethyl gallate (EG) and propyl gallate (PG), had no antiviral activity in 10 min exposure (Table 1 and Fig. 6), even though they were cytostatic (EG) or cytotoxic (PG) after 48 or 72 h exposure times to 100 µM. The gallate esters of longer chain fatty acids, lauryl- and octyl-gallates, had potent and broad spectrum activity when virions were exposed to them for 10 minutes (Fig. 6), although they also increased cell doubling times by more than 1.5-fold during 48 and 72 h exposure times to just 10 µM. There were no differences in the antiviral potencies of octylgalloyl (OG) or laurylgalloyl (LG).

**Fig. 6.**
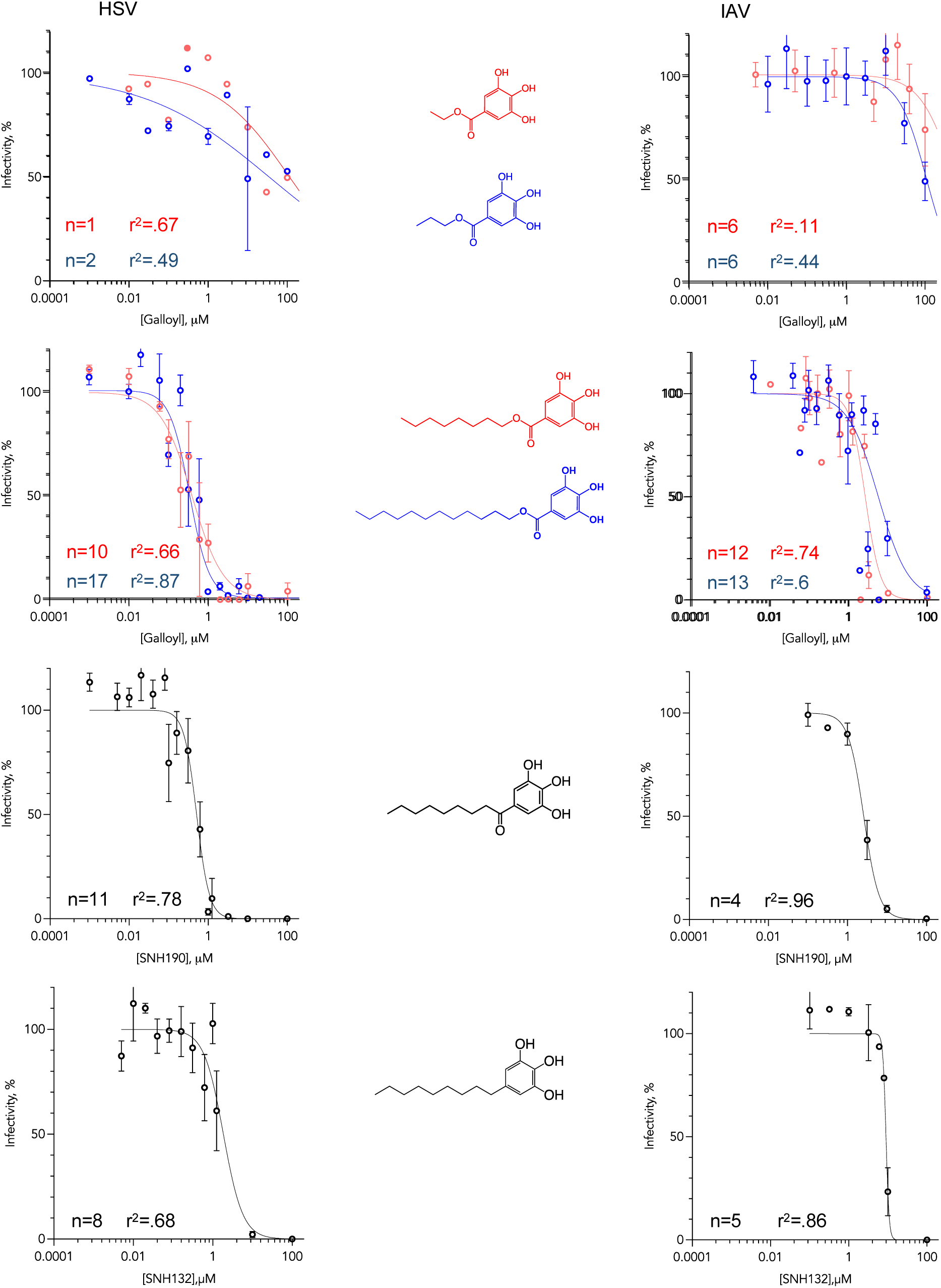
Effects of removing the ester linkers in alkyl derivatives. Dose-response line graphs showing the inhibition of infectivity of HSV-1 or IAV by the indicated alkyl polyphenol derivatives. HSV-1 or IAV virions were mixed with test compounds incubated at 37°C for 10 minutes before infecting Vero (HSV-1) or MDCK (IAV) cells with 200 PFU. Average ± SEM, n= 1 to 12 biological independent experiments.

We propose that the long chain gallates likely aggregate in acqueous environment through their hydrophobic moieties, effectively resulting in polygallates. Regardless of the specific mechanism, we synthesized a series of compounds in which the ester group was disrupted. Replacement of the ester with a keto group resulted in no change in potency or broad spectrum activity (Table 1 and Fig. 6, SNH190 vs OG/LG), or cytotostatic effects after 48 and 72 h exposure. Removal of the keto group decreased potency by about 3.5-fold (Table 1 and Fig. 6, SNH132 vs OG/LG) while also decresing the cytostatic effects, in that no effects on cell doubling time were observed after 48 or 72 h exposure to 100 µM.

### Selection for IAV resistance against EGCG

We had biased the screens towards compounds that act on virions, not cells, and therefore we pursued selection for resistance to confirm a viral target. As we proposed that the synthetic compounds act via the same mechanism of action as EGCG, we used commercially available EGCG for this purpose. We performed parallel selection using two different IAV strains, PR8 (H1N1) and Aichi (H3N2), following two variations of a selection protocol that we have used before (29, 98). For PR8, we started at lower concentration, increasing it in each passage or keeping it constant to allow for titres to recover. For Aichi, we started at a higher concentration, although we had to lower it after passage 3 for the titres to recover. Serial passage in increasing EGCG concentrations promptly selected for resistance in both strains. The titres in the preseence of EGCG typically decreased with respect to those in DMSO in each passage when the EGCG concentration was increased, but promtply recovered to similar titres as in DMSO. After 5 passages, both PR8 and Aichi grew to the same titers in the presence of 120 or 0 µM EGCG (Fig. 7); we performed one additional passage and plaque purified the resistant strains.

**Fig. 7.**
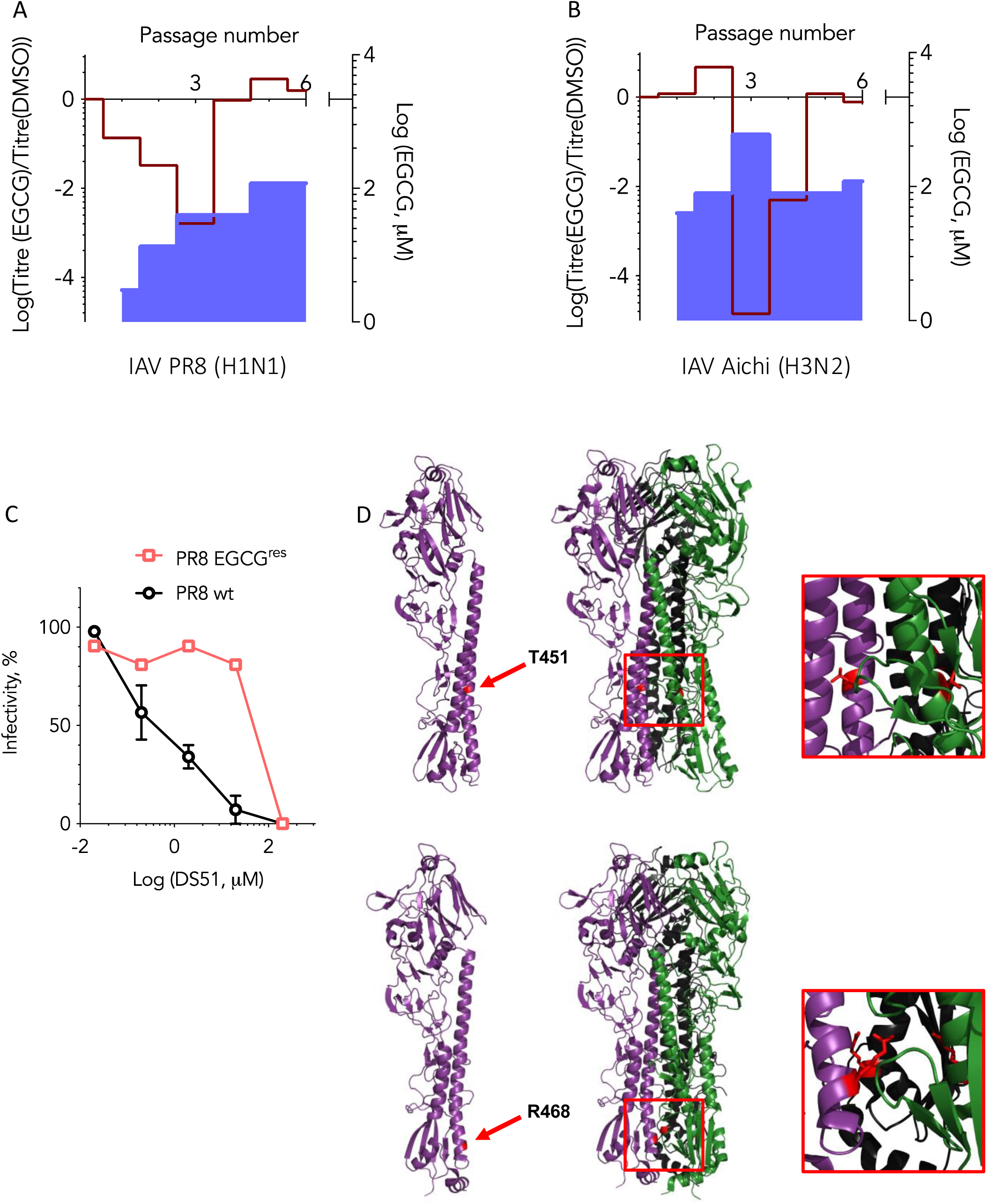
Selection for resistance to EGCG by two strains of IAV. A and B, Line graphs presenting the ratio in viral titres per cell during passages (red line) in the presence of increasing concentrations of EGCG (blue surface) or in DMSO. The titres produced during each passage in the presence of EGCG were divided by the titres of the corresponding passage in DMSO; the x-axis at 1 indicates equal titres in the presence or absence of EGCG. A, PR8 (H1N1); B, Aichi (H3N2). **C,** dose response inhibition of infectivity of wild-type or a EGCG^r^ strain of PR8 in the presence of DS51. **D and E,** mapping the mutations selected for in all plaques of Aichi (D), or PR8 (E) to the stalk of HA.

A PR8 EGCG^r^ strain was cross-resistant to the trigalloyl DS51 (66-fold EC_50_ increase), indicating a common mechanism of resistance, and further confirming that the observed DS51 antiviral effects were not a result of cytotoxicity or cytostatic effects. We sequenced the HA genes in different plaques of both isolates. In both PR8 and Aichi, we identified a single nonconservative mutation in all plaques. In both cases the mutation replaced a charged or polar amino-acid in the inside of the base of the stalk in the pre-fusion neutral pH conformation (T434 in PR8 or R453 in Aichi) (Fig. 7) by a non-polar amino-acid (I in PR8 or G in Aichi). No mutations were identified in the strains passaged in DMSO as a control.

### Selection for HSV-1 resistance against EGCG

We also selected for HSV-1 resistance against EGCG. Like for IAV, resistance was promptly selected for and the titres produced in the presence of EGCG at passage 8 had recovered to the initial titres (Fig. 8). During this selection, we noted that the percentage of syncitial plaques increased with the number of passages. After 8 passages, half of the plaques were syncitial (Fig. 8), fully consistent with the phenotype of the plaques obtained during selection for heparin resistance (99) and indicating that the selection had resulted in an enrichment in variants that spread preferentially by direct cell-to-cell spread. *syn*^+^ mutations map to six different HSV-1 proteins, including gC and the cytoplasmic tail of gB (99, 100), the two HSV-1 glycoproteins that bind to GAG (99, 101). Syncitial mutations are consistent with the proposed mechanism of inhibition, as they bypass the inhibited step and have been previoulsy identified in selection for resistance to heparin, but do not necessarily point to a potential binding site. We therefore did not sequence these mutants.

**Fig. 8.**
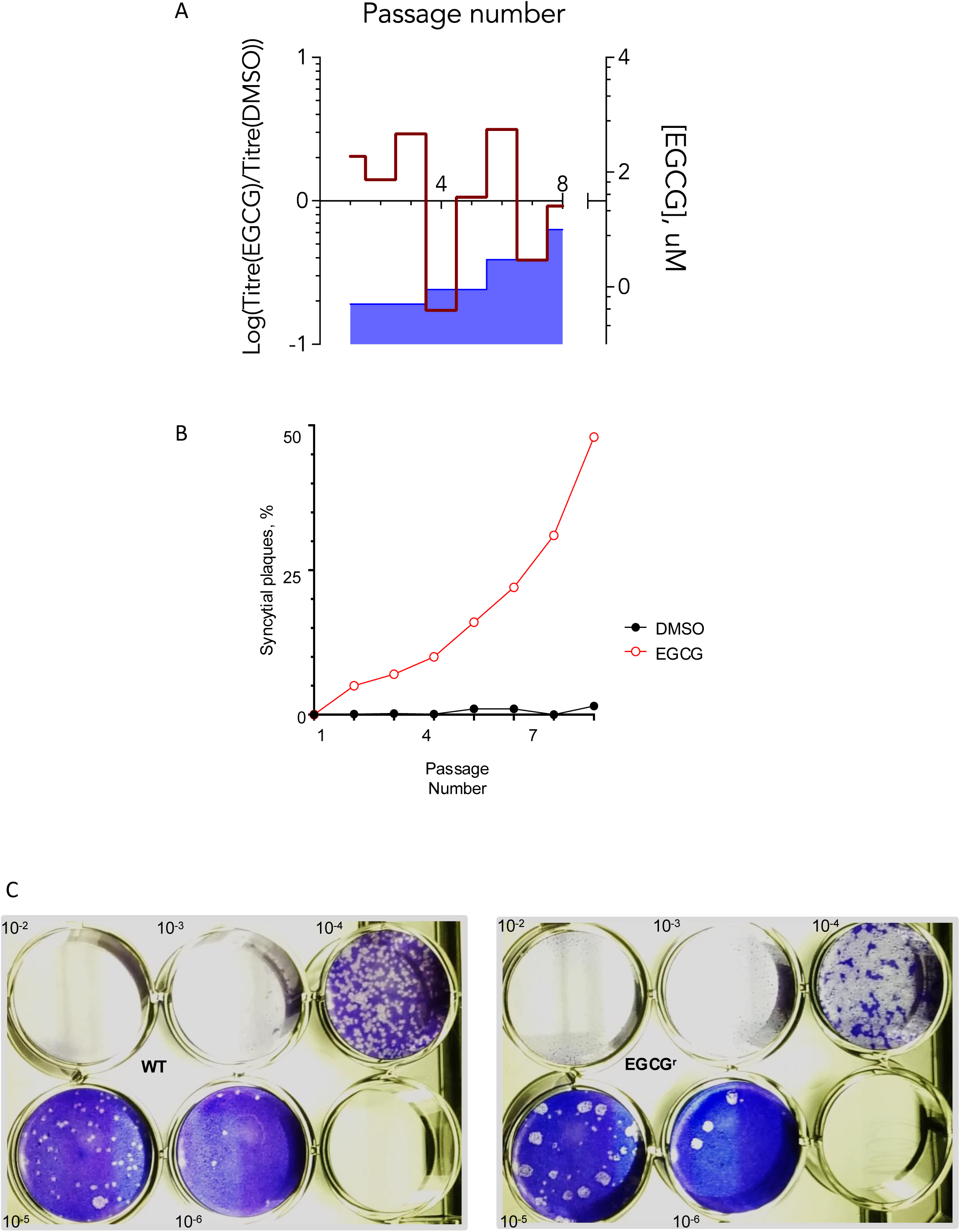
Selection for resistance to EGCG by HSV-1. **A**, Line graphs presenting the ratio in viral titres per cell during passages (red line) in the presence of increasing concentrations of EGCG (blue surface) or DMSO. The titres produced during each passage in the presence of EGCG were divided by the titres of the corresponding passage in DMSO; the x-axis at 1 indicates equal titres in the presence or absence of EGCG. **B,** percentage of *syn*+ plaques after each passage in the presence of increasing concentrations of EGCG. **C,** photographs of plaque morphology (both plates infected with equivalent number of PFU from passage 8 in the presence or absence of EGCG, fixed and stained at the same times).

### The galloyl compounds are broad spectrum antivirals that inhibit human coronaviruses that attach to SG or GAG

Based on the previous results, we proposed that these polygalloyl compounds have broad spectrum antiviral activity by targeting the virions to inhibit their attachment to cells. However, we had tested them against only one virus that attaches to GAG, and one that attaches to SG, HSV-1 and IAV, respectively, and these viruses are unrelated to each other. They differ in their glycan attachement but also their mechanisms of entry and replication. Moreover, we had tested their effects only on virion infectivity (i.e., virucidal test), to explore the chemical space without the complexities of potentially including compounds acting on virions, cells, or both, as well as to include compounds that were cytotoxic upon long exposures. We thus tested the potential broad spectrum antiviral activity of the selected compounds on viruses that attach to SG or GAG and are unrelated to IAV or HSV-1, but related to each other in general entry and replication mechanisms. To this end, we used the coronaviruses hCoV OC43 or SARS-CoV-2.

We first performed a primary CPE screen using human coronavirus hCoV OC43. hCoV OC43 attaches to SG (48) and most compounds had lower potency against influenza than HSV-1; hCoV OC43 was therefore expected to be more stringent for the screens. The screen was developed to resolve between inhibitors of spread or replication. In brief, susceptible cells infected at low (0.3) or high (3) multiplicity of infection (moi), or mock infected cells, were treated with 10 µM test compounds after removing the inoculum. Two and three days after infection, cell viability was quantitatively evaluated by CellTiter-Glo. Uninfected and untreated cells and infected untreated cells indicated relative 100 and 0% viability, respectively (Fig. 9). We set the threshold at 70% relative viability at 2 and 3 days post infection, to minimize spurious hits. Of the 32 preselected gallate compounds and 10 positive controls screened, 25 gallate compounds and 9 controls scored as positive or potentially positive at low moi and none at high moi (Fig. 9), which is consistent with the proposed mechanism of action.

**Fig. 9.**
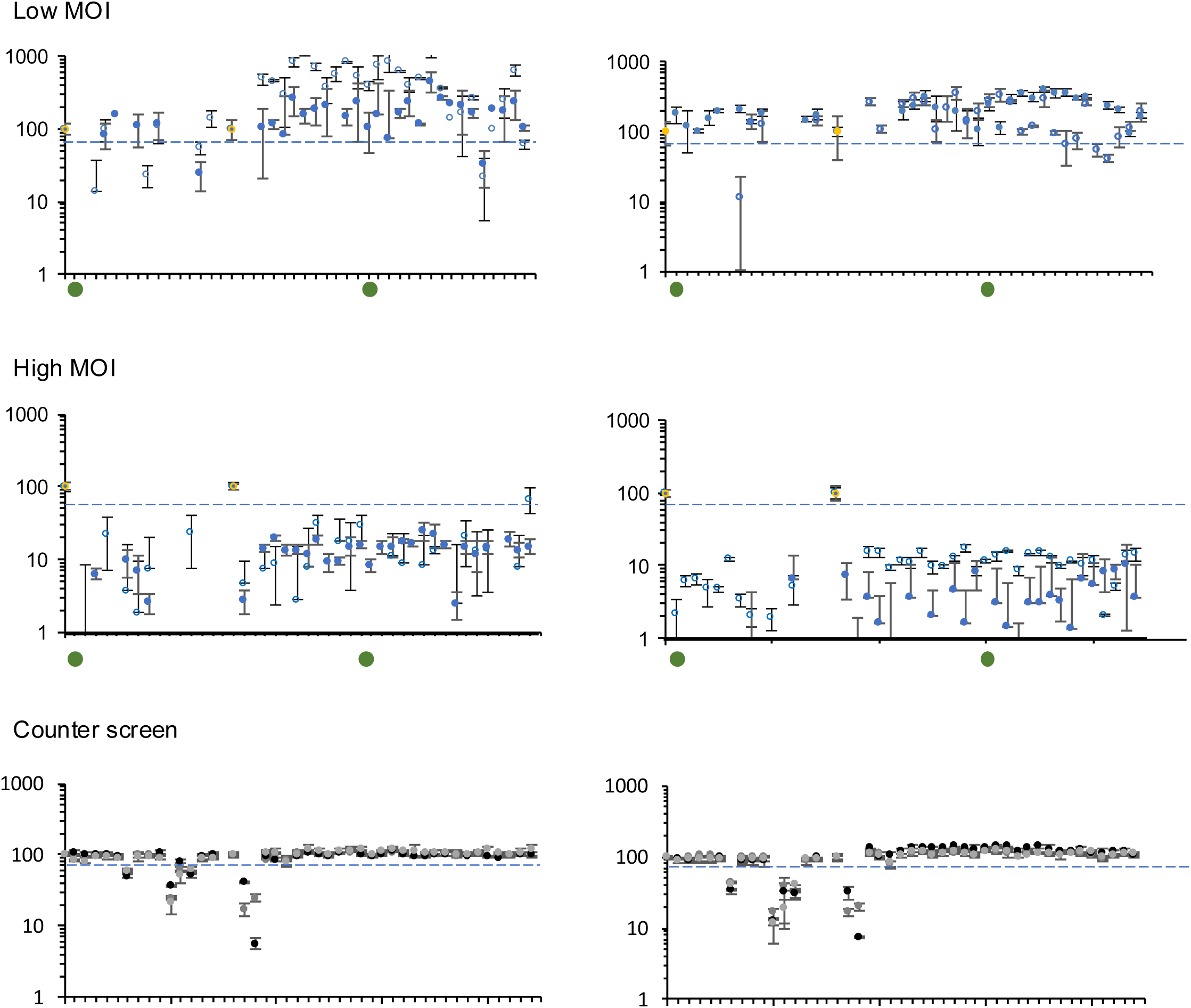
Screening for inhibitors of OC43 infection spread. Vero E76 **c**ells were infected at low (0.3 FFU/cell) or high (3 FFU/cell) multiplicity with OC43 or left uninfected. Inocula were removed 1 h later and infected or mock-infected cells were overlaid with medium containing 10 µM of test compounds in triplicates. Cell viability was evaluated 2 (left panels) and 3 (right panels) days later, setting he viability of infected untreated cells as 0% and that of uninfected untreated cells as 100%. Compounds that resulted in viability of infected cells of 70% of that of uninfected untreated cells or higher at low but not high moi were considered hits. As expected from inhibitors of infectivity, no compound scored as a hit at high moi, when all cells were infected before adding test compounds. Compounds that inhibited viability to below 70% of that of untreated and uninfected cells were considered cytostatic or toxic at 10 µM. Blue circles, viability percentage of infected treated cells (test compounds); yellow circles, uninfected-untreated cells (100% viability); green circles, infected non treated cells (0% viability); blue dashed line, 70% viability (cut off); grey and black circles, not infected treated cells. Individual results of biological replicates are presented (left and right plots) as the average ± SD of the technical replicates in each repeat; n=2 or 3 biological independent experiments.

We next validated selected hits by dose-response analyses. EGCG inhibited the replication of hCoV OC43 in three days by about 20-fold, with an EC_50_ of about 8 µM (Table 1 and Fig. 10). The digallates of flexible linkers were only weakly active or inactive in this model. SNH32 had no antiviral activity, whereas SNH34 and SNH36 had potencies in the mid to high micromolar range. Oxygen atoms in the linker, or the replacement of the ester by amide groups resulted in similar weak activity. The replacement of the flexible linkers by rigid non-planar ones also resulted in mostly weak activity in cell culture, which was not significantly affected by the position of the galloyl substituents (Table 1 and Fig. 10). It is possible that all these two ester-containing compounds are unstable during the three days of treatment in tissue culture, while still scoring as positive in the CPE screens by inhibiting replication for just long enough to prevent the virus from killing more than 30% of cells in the 2 and 3 days evaluated.

**Fig. 10.**
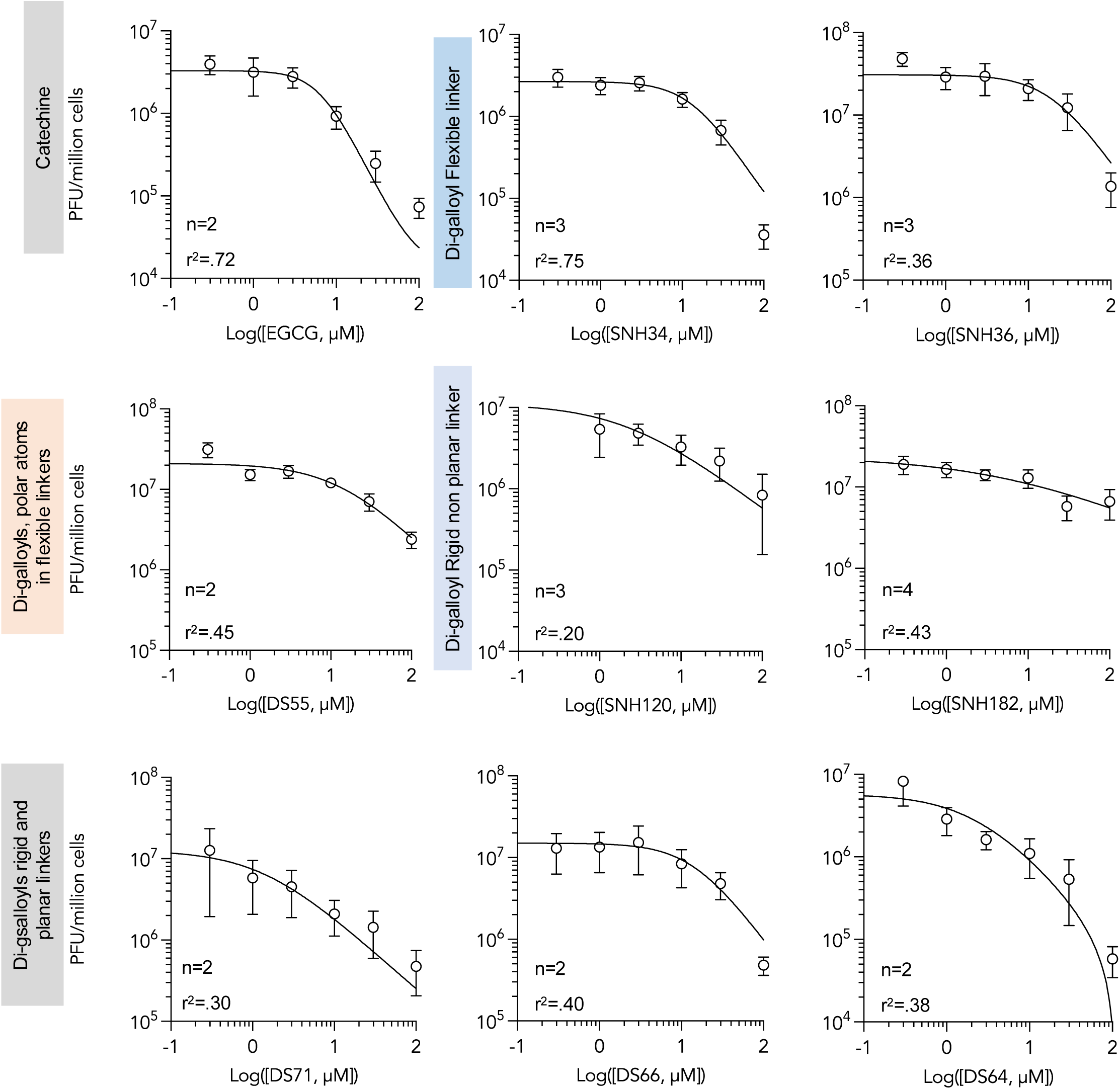
Dose response effects of selected gallates on OC43. Dose-response line graphs showing the inhibition of production of infectious virus by the indicated polygalates in cells infected at low moi before treatment. Monolayers were infected with OC43 at an moi of 0.05 FFU/cell; inoculum was removed 1 h later and infected cells were overlaid with medium containing the indicated concentrations of test compound. Supernatants were harvested 72 h later and viral in the supernatants was titrated by foci formation via immunofluorescence. Average ± SEM, n= 2 to 4 biological independent experiments.

The digallates of rigid and planar linkers had some antiviral activity agasint OC43, but the activities of DS71 and DS64 were observed at cytostatic or close to cytostatic concentrations only and were not analyzed in further detail. The non-cytostatic DS66 had similar activity at high concentrations only (Table 1 and Fig. 10).

In contrast, the trigallates DS74 and DS51 and the tetragalloyl SNH78 inhibited the replication of hCoV OC43 by about 100-fold with EC_50_ below 10 µM (Table 1 and Fig. 11). Although DS51 is cytostatic at 100 µM (not at 10 µM), the non-cytostatic DS74 and SNH78 had similar potencies, suggesting that cytostaticity and antiviral activity are different activities of these compounds. As for HSV-1 or IAV, attemtps to test higher concentrations were unsuccesful.

**Fig. 11.**
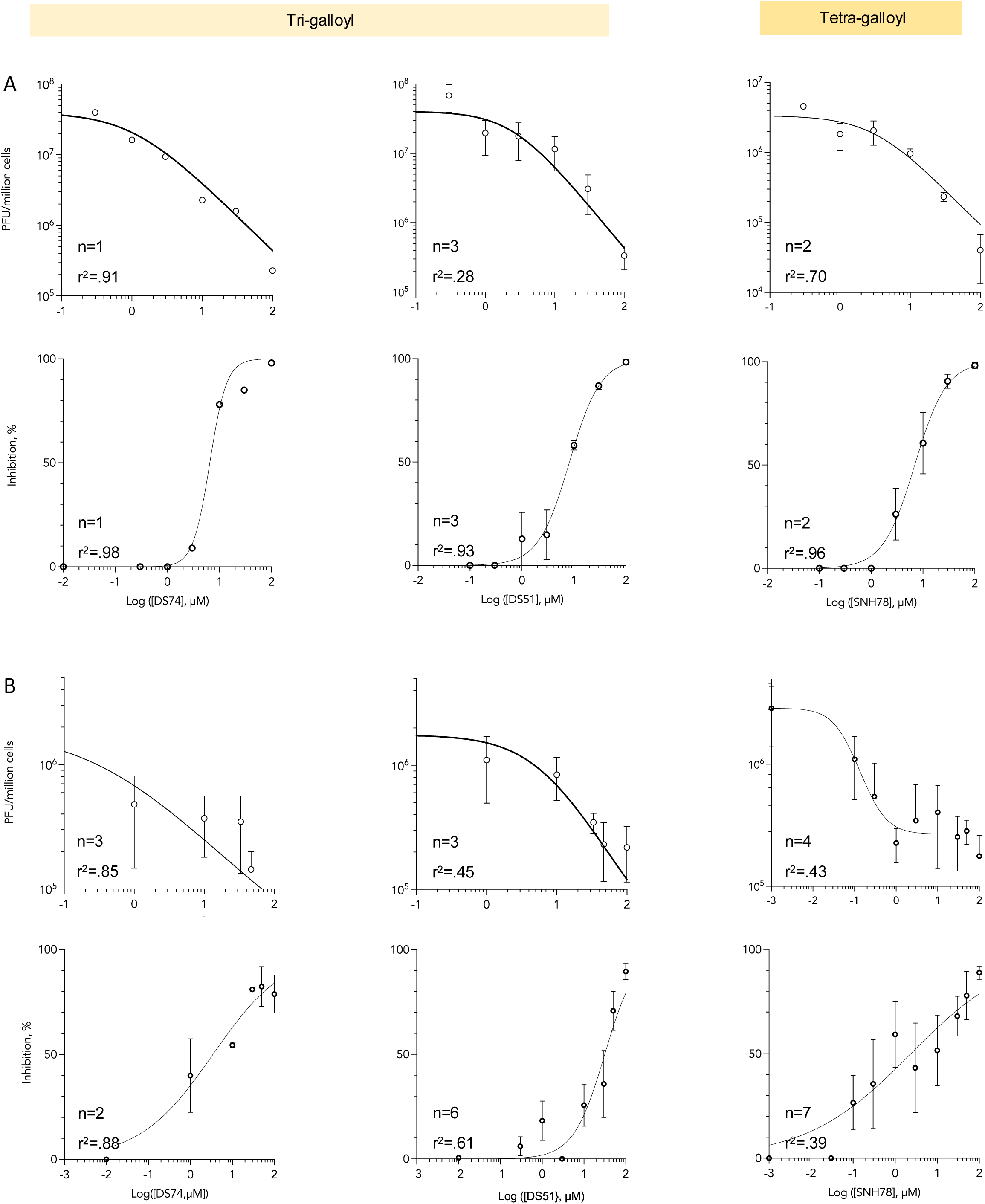
Dose response effects of DS51 and SNH78 on hCoV OC43 and SARS-CoV-2. Dose-response line graphs showing the inhibition of production of infectious virus by the indicated polygalates in cells infected at low moi before treatment. **(A)** Monolayers were infected with OC43 at an moi of 0.05 FFU/cell; inoculum was removed 1 h later and infected cells were overlaid with medium containing the indicated concentrations of test compound. Supernatants were harvested 72 h later and viral in the supernatants was titrated by foci formation via immunofluorescence. **(B)** Monolayers were infected with SARS-CoV-2 at an moi of 0.03 FFU/cell; inoculum was removed 1 h later and infected cells were overlaid with medium containing the indicated concentrations of test compound. Supernatants were harvested 48 or 72 h later and virus in the supernatants was titrated by plaque assays. Average ± SEM, n= 1 to 5 biological independent experiments.

Consistent with their effects on hCoV OC43, digalloyl esters of flexible or rigid-non planar linker, were inactive against SARS-CoV-2. The digalloyl ester of the rigid and planar linker DS71 had only weak activity, at cytostatic concentrations. DS51 was also weakly active against SARS-CoV-2, at concentrations close to cytostatic (EC_50_ 5.3 µM), whereas SNH78 was more potent, inhibiting replication by approximately 16-fold at 1 µM with an EC_50_ of 2.3 µM (Table 1 and Fig. 11). Once again, attempts to test higher concentrations were unsuccessful.

## Discussion

Considering that attachment to SG and GAG use different mechanisms, it is unsurprising that no compound had been identified to inhibit attachment to both types of glycans until we and others unexpectedly found that the green tea catechin EGCG does so (67). Although EGCG has vastly different potencies against HSV-1 and IAV, these findings prompted us to explore the possibility of identifying new chemical entities that could inhibit attachment to both types of glycans with similar potency.

Previous attempts to modify catechins used oxidation-induced dimerization (102) or lipid conjugations (103). These approaches create more complex derivatives of the natural compounds; in contrast we chose to deconstruct the base molecule, then design and synthesize new compounds containing the moieties predicted to be necessary for antiviral activity and omitting those not expected to be important. We thus produced simpler synthetic compounds that define a minimal chemical scaffold with broad spectrum antiviral activity.

EGCG and related catechins inhibit HSV-1 and 2 attachment to GAG, target the GAG-binding gB glycoprotein, and inhibit hemagglutination of IAV (67, 102, 104). Here, we show that catechins and digallate compounds also inhibit the spread of human coronavirus hCoV OC43 when added to cells already infected at low moi, but not its replication in cells already infected at a high moi. Therefore, we favor the model in which the synthetic gallates operate by the same model of inhibiting attachment to cellular glycans, although we have not formally explored their mechanism of action against coronaviruses. These conclusions are also supported too by the results obtained testing the effects of EGCG on GAG attachment by hCoV OC43 and SARScCoV-2 S, the effects of EGCG and marine sulfated glycans on pseudotyped SARS-Cov-2 attachment and entry (51, 62), and of EGCG on hCoV OC43 (51). We show here that the compounds are also active against SARS-CoV-2, an emerging coronavirus that attaches to GAG, in infections at low moi, supporting the activity of these compounds against unrelated viruses that attach to SG or GAG, IAV and OC43 or HSV-1 and SARS-CoV-2, respectively.

In our first attempt to identify the specific target and mechanism of action, we selected for resistance against EGCG. The mutants selected for in IAV PR8 and Aichi both map to the base of the HA stalk. Mutations in the non surface exposed alanine 388 to threonine or valine in the base of the HA stalk confer resistance to several neutralizing monoclonal antibodies (105-107) without disrupting the antibodies binding site (the mutation is not exposed). Similar phenotype results from mutations at asparagine 391 to tyrosine or glycine (108). Stalk mutations that destibilize HA also confer resistance to arbidol (109). The mutations we selected for may thus have conferred resistance by minimizing the dependence on receptor binding to trigger the conformation changes in HA required for fusion. Alternatively, as the HA stalk is the binding site for small molecules that inhibit IAV fusion (110), it may also be the binding site for the polygalloyl compounds presented here.

The HSV-1 ECGC resistant mutants were syncitia inducing (*syn*+), consistently with previous results selecting for resistance agains heparin (99). Mutations in six loci, including the cytoplasmic tail of the GAG- and EGCG-binding gB, result in the direct cell-to-cell fusion that produces the syncytia (99, 100, 111, 112). These mutations bypass the step at which virion glycan attachment plays its role and thus this mechanism of escape needs not affect compound binding. We therefore decided against mapping these mutations, as they would not be informative for our purposes. In summary, the selection for resistance in IAV or HSV-1 may have not been informative with respect to the specific binding sites of these comounds but it is fully consistent with the proposed mechanism of action by inhibiting attachment.

Attempts to evaluate higher concentrations often resulted in high variability and inconsistent dilution effects, suggesting aggregation. We thus limited the analyses to lower concentrations, at the cost of not being able to identify the potential maximum extent of inhibition.

The specific chemical entities used were selected to most efficiently test the hypothesis that it is possible to inhibit attachment of unrelated viruses to different glycans with similar potency. Although these molecules are not pharmacologically optimized, having ester links and polyphenols, and are limited by solubility, they chart a path toward development of new chemical entities acting by this mechanism of action.

In conclusion, we show here that it is possible to develop small molecule synthetic inhibitors of attachment to GAG and SG. Even if this approach were to produce molecules with mild potency only, this type of activity would still be highly desirable as a first-line countermeasure for any unknown emerging virus, to contain the spread sufficiently until more potent specific antivirals are eventually developed.

## Materials and Methods

### Compounds

The detailed procedures for the synthesis of all compounds are described in the supplementary information. Test compounds were stored at -20°C as 100 mM DMSO stock solutions in aliquots. Each aliquot was thawed no more than 3 times.

### Cells and viruses

Vero African green monkey kidney cells (ATCC, CRL-1587, CRL-1586), MDCK (ATCC CCL-34), and human colorectal cancer cells HRT18G (CRL-11663) were maintained in DMEM supplemented with 5 % FBS at 37°C in 5% CO_2_.

Stocks of HSV-1, strain KOS p5 after plaque purification of the original ATCC stock (VR-1493) to minimize the presence of *syn*^+^ plaques, were grown in Vero cells. In brief, cells were infected at an moi of 0.01 PFU/cell. Inoculum was removed 1h later and infected cells were overlaid with DMEM-5% FBS. Cells were harvested by scraping when the monolayers showed 4+ CPE, typically about 4 days after infection. Cells and supernatant were fractionated by centrifugation. The cell pellet was resuspended with 300 µl of DMEM, freeze thawed three times, sonicated, and clarified by centrifugation to remove cell debris; the cell pellet supernatant was then kept on ice till use. The supernatant was pelleted at 10,000 g for 2 h and the virion pellet was resuspended in 300µl of the cell supernatant. The resulting viral stock was aliquoted and stored at -80⁰C. Virus stocks were titrated by plaque assay (1.4x10^9^ PFU/ml).

Stocks of IAV (originally obtained from Dr. Veronica von Messling, German Federal Ministry of Education) (67) were prepared by infecting MDCK cells at an moi of 0.01 PFU/cell. Inoculum was removed after 1 h, cells were washed, and fresh DMEM supplemented with 0.2% BSA and 2µg/ml TPCK trypsin. Supernatant was harvested when 99% of cells had obvious CPE and clarified by centrifugation. Virions were pelleted at 10,000g at 4°C for 2 h, resuspended in DMEM, titrated by plaque assay (1.4x10^7^ PFU/ml), aliquoted and frozen at -80°C until use.

Stocks of hCoV OC43 (a generous gift from Dr. D. Diel, originally from ATCC, VR-1740) were prepared by infecting HRT-18G or Vero cells at a multiplicity of infection of 0.01 PFU/cell for 1 h at 37°C. Inocula were removed and cells were washed twice with cold DMEM. Fresh pre-warmed DMEM – 5 % FBS was added to the infected cells, which were then incubated at 37°C in 5 % CO_2_ and harvested when full CPE was visible (typically, 4 to 5 days after infection). Viral stocks were harvested in DMEM with no FBS, titrated by immunofluorescence (3x10^7^ FFU/ml), and aliquoted and frozen at -80°C until use.

Stocks of SARS-CoV-2 USA-WA1/2020 were grown from BEI stock NR-52281 and used at passage 1 or 2 only. In brief, Vero cells were infected at MOI of 0.002 PFU/cell for 1 h at 37°C. Inoculum was then removed, fresh pre-warmed DMEM – 5 % FBS was added to and cells were incubated at 37°C in 5 % CO_2_ and harvested at 48 h post infection. Collected supernatants were cleared by centrifugation, aliquoted, and frozen at -70°C until use. Stocks (1.1x10^6^ PFU/ml) were titrated on Vero E6 cells by plaque assay.

### Cell viability and cell doubling

To evaluate the effects of test compounds on cell viability and doubling, 2.5 x 10^3^ Vero cells were seeded into each well of 96 well plates and incubated at 37 °C in 5% CO_2_ for 18 h. Semi logarithmic serial dilutions of compounds and vehicle control in DMEM-5% FBS were prepared just before use in a 96 deep well plate. Cells were washed twice with warm DMEM before 100 µL compound solution was transferred with a multichannel pipette in technical triplicates. Cells were incubated with compounds for 72 h at 37°C evaluating cell viability daily.

### Virion infectivity assays

To test the effects of the test compounds on the infectivity of HSV-1 virions, Vero 76 cells were plated at 1.5x10^6^ per well in 12 well plates. HSV-1 was diluted 20 h later to twice the target concentration of 200 pfu per 100 µl and logarithmic or semi-logarithmic test compound dilutions were prepared at 2X final concentration in complete medium. Virions were treated with test compound by mixing 1:1 ratio of diluted virus and diluted compounds, both pre-warmed to 37 °C, and incubating the mix for 10 minutes in 37°C water bath. Treated virions were then transferred to ice bath until infection. Cells were washed once with cold serum free DMEM and inoculated with 100 µl of treated virions. After adsorption for 1 h in 5% CO_2_ at 37°C, inocula were removed and cells were washed twice with cold DMEM. Afterwards, infected cell monolayers were overlaid with 2% methylcellulose in DMEM supplemented with 5% FBS in and incubated in 5% CO_2_ at 37°C until plaques were visible and well defined, typically 48-72 h. Cells were then fixed and stained with 2% crystal violet in 17% methanol for 18-24 h. Wells were washed with tap water, plaques were counted, and numbers were plotted, graphed and statistically analyzed using GraphPad Prism 9.0.

To test the effects of the test compounds on the infectivity of IAV virions, MDCK cells were plated at 0.9x10^6^ per well in a 6 well plates (about 2 plates per compound). Virion treatment and adsorption was performed as for HSV-1, and then infected cell monolayers were overlaid with 0.8% Agarose + 1µg trypsin-TPCK and incubated in 5% CO_2_ at 37°C until plaques became visible and well defined, typically in about 36 h. Cells were fixed, stained and washed, and plaques were counted and analyzed as for HSV-1.

### hCoV OC43 CPE screening

The potential antiviral activity of selected test compounds against hCoV OC43 was first evaluated by cell viability screening. Vero cells (2.5 x 10^3^/ well) were seeded in 96 well plates and incubated at 37°C in 5% CO_2_ for 18 h before infecting them with hCoV OC43 at an MOI of 0.3 or 3 FFU/cell for 1 h at 37 °C. Inocula were removed and cells were washed twice with cold DMEM. Test compounds prepared at 10 µM in prewarmed DMEM 5 % FBS were added to infected and uninfected cells, which were then incubated at 37°C in 5 % CO_2_. Non infected (100% viability) and infected (0% viability) untreated controls were included in each plate. Cells were incubated with compounds for up to 72 h at 37°C. Medium was removed and cells were washed twice with phenol red free DMEM at 37°C. Plates were cooled down to room temperature, medium was removed, and 50 µL phenol red free DMEM and 50 µL Cell-titer-glo Luminescent Cell viability assay (Promega, G7571/2/3) were added and mixed in an orbital shaker for 2 min. Afterwards, 80 µL of the mix was transferred onto white 96 well plates and incubated for 10 min at RT for signal stabilization. Luminescence was read in the Glomax Multidetection System (Promega) with an integration time of 0.5 sec.

Means of technical triplicates of test samples were normalized to the means of the controls. Compounds were considered hits when the viability of treated and infected cells was at or above 70% of that of the uninfected and untreated cells (0 %, viability of infected and untreated cells and 100 % viability of uninfected untreated cells). A counterscreen with uninfected cells was performed to identify cytotoxic compounds, using parallel uninfected plates.

### Dose-response analyses of hCoV OC43 replication

7.5x10^4^ Vero cells were seeded into each well of 24 well plates and incubated at 37°C in 5 % CO_2_ for 18 h before infecting with OC43 at an moi of 0.05 FFU/cell for 1 h at 37 °C. Inocula were removed and cells were washed twice with cold DMEM. Test compound dilutions in prewarmed DMEM 5 % FBS was added to the infected cells, which were then incubated at 37°C in 5 % CO_2_ and the supernatant was harvested at 72hpi. Supernatants were titrated by focus-forming assay on Vero cells. Ten-fold serial dilutions of the harvested supernatants were prepared in DMEM. Vero cells (7.5 × 10^4^ cells/well in 24-well plates) were then infected with 75 µL of the appropriate dilution for one h. Then, monolayers were overlaid with 2 % methylcellulose and 5 % FBS and incubated at 37°C for 72 h. Overlaid cells were washed three times with warm PBS and cells were fixed with buffered formalin for 10 min at 4°C. Formalin was removed and cells washed three times with cold PBS, before permeabilizing with 0.5% triton for 13 min at RT and washing three times with PBS. Fixed cell monolayers were blocked with 5% goat serum and 2% BSA at RT for 1 h, or overnight at 4 °C, and incubated with primary mouse IgG anti-HCoV-OC43 antibody (Chemicon, MAB9012) diluted 1:1000 in 1% BSA for 4 h at RT or overnight at 4 °C. After primary antibody, cell monolayers were washed three times with PBS before incubating with secondary goat anti-mouse IgG Alexa Fluor 488 antibody (Nikon Eclipse Ts2R) for 1 h at RT. Monolayers were finally washed with PBS and foci of infected cells were counted using a fluorescence microscope (Invitrogen, A11001). Means of results were normalized to the means of the vehicle controls, and then analyzed by nonlinear regression dose responses, and graphed, using GraphPad Prism 9.0. Two to four independent biological replicates were performed for each treatment (n=2 to 4).

### Dose-response analyses of SARS-CoV-2 replication

All procedures using live SARS-CoV-2 were performed in the BSL3 suite following approved BSL3 protocols. In brief, Vero cells were seeded at 1.5x10^5^, 7.5x10^5^, or 2.5x10^3^ cells/well in 12, 24, or 96 well plates, respectively, washed twice with cold DMEM, and infected at an MOI of 0.003 or 0.03 PFU/cell for 1 h in 5 % CO_2_ at 37°C, rocking and rotating every 10 minutes. Inocula were removed, cells were washed twice, and test compound dilutions of in prewarmed DMEM supplemented with 5% FBS were added to the infected cells and incubated at 37°C. Forty-eight to seventy two hours after infection, depending on well and moi, 150 µL or 50 µl of supernatant was transferred into a 1.5 mL screw cap tube, and cleared at 1100 rpm for 5 minutes at 4°C. Cleared supernatant was transferred into a new tube, quickly frozen and stored at -80°C until titrating. Alternatively, supernatant was transferred into a round bottom 96 well plate, quickly frozen, and stored at - 80°C until titrating.

To titrate replicated virus, Vero E6 cell monolayers were seeded at 1.0x10^5^ cells/well in 24 well plates and infected with ten-fold dilutions of test sample in DMEM for 1 h in 5 % CO_2_ at 37°C, rocking and rotating every 10 minutes. Cell monolayers were then overlaid with 2 % methylcellulose (Sigma-Aldrich M0387-250G) in DMEM supplemented with 5 % FBS and incubated in 5 % CO_2_ at 37°C until well-defined plaques developed (typically 3 to 4 days). Monolayers were then fixed with 4 % formalin (Sigma-Aldrich, 1.04003.1000) for 20 minutes at room temperature (RT) and stained overnight with crystal violet (Sigma-Aldrich, 0528-100G) in 17 % methanol before washing and counting plaques.

